# Area of Habitat maps for the world’s terrestrial birds and mammals

**DOI:** 10.1101/2022.05.13.489640

**Authors:** Maria Lumbierres, Prabhat Raj Dahal, Carmen D. Soria, Moreno Di Marco, Stuart H. M. Butchart, Paul F. Donald, Carlo Rondinini

## Abstract

Area of Habitat (AOH) is ‘the habitat available to a species, that is, habitat within its range’. It complements a geographic range map for a species by showing potential occupancy and reducing commission errors. AOH maps are produced by subtracting areas considered unsuitable for the species from their range map, using information on each species’ associations with habitat and elevation. We present AOH maps for 5,481 terrestrial mammal and 10,651 terrestrial bird species (including 1,816 migratory bird species for which we present separate maps for the resident, breeding and non-breeding areas). Our maps have a resolution of 100 m. On average, AOH covered 66±28% of the range maps for mammals and 64±27% for birds. The AOH maps were validated independently, following a novel two-step methodology: a modelling approach to identify outliers and a species-level approach based on point localities. We used AOH maps to produce global maps of the species richness of mammals, birds, globally threatened mammals and globally threatened birds.

## Background & Summary

Knowing the distribution of species is crucial for effective conservation action. However, accurate and high-resolution spatial data are only available for a limited number of species^1,2^. For mammals and birds, the most comprehensive and widely used global distribution dataset is the set of range maps compiled as part of the assessments for the International Union for Conservation of Nature (IUCN) Red List. These represent each species’ distributional limits and tend to minimise omission errors (i.e. false absences) at the expense of commission errors (i.e. false presences)^3,4^. Therefore, they often contain sizeable areas that are not occupied by the species.

Maps of the Area of Habitat (AOH; previously known as Extent of Suitable Habitat, ESH) complement range maps by indicating potential occupancy within the range, thereby reducing commission errors^5^. AOH is defined as ‘the habitat available to a species, that is, habitat within its range’^5^. These models are produced by subtracting areas unsuitable for the species within their range, using information on each species’ associations with habitat and elevation^5–8^. Comprehensive sets of AOH maps have been produced in the past for mammals^6^ and amphibians^7^, as well as subsets of birds^8,9^. The percentage of a species’ range covered by the AOH varies depending on the methodology used to associate species to their habitats, and their habitats to land-cover, the coarseness of the range map, the region in which the species is distributed, and the species’ habitat specialisation and elevation limits^5^. For example, Rondinini et al.^6^ found that, when considering elevation and land cover features for terrestrial mammals, the AOH comprised on average 55% of the range. Ficetola et al.^7^ obtained a similar percentage when analysing amphibians (55% for forest species, 42% open habitat species and 61% for habitat generalists). Beresford et al.^8^ found that AOH covered a mean of 27.6% of the range maps of 157 threatened African bird species. In 2019, Brooks et al.^5^ proposed a formal definition and standardised methodology to produce AOH, limiting the inputs to habitat preferences, elevation limits, and geographical range.

AOH production requires knowledge on which habitats a species occurs in and their location within its range^1^. Information on habitat preference is documented for each assessed species in the IUCN Red List^10^, following the IUCN Habitats Classification Scheme^11^. However, IUCN does not define habitat classes in a spatially explicit way, therefore, we used a recently published translation table that associates IUCN Habitat Classification Scheme classes with land cover classes^12^. Species’ elevation limits were also extracted from the IUCN Red List.

We developed AOH maps for 5,481 terrestrial mammal and 10,651 terrestrial bird species. For 1,816 bird species defined by BirdLife International as migratory, we developed three AOH maps, one for the resident range, one for the breeding range and one for the non-breeding range. The maps are presented in a regular latitude/longitude grid with an approximate 100m resolution at the equator. On average, the AOH covers 66 of the geographical range for mammals and 64+27% for birds. We used the resulting AOH maps to produce four global species richness layers: for mammals, birds, globally threatened mammals and globally threatened birds.

The AOH maps have multiple conservation applications^5,13,14^, such as assessing species’ distributions and extinction risk, improving the accuracy of conservation planning, monitoring habitat loss and fragmentation, and guiding conservation actions. AOH has been proposed as an additional spatial metric to be documented in the Red List^5^, and is used as an assessment parameter in the identification of Key Biodiversity Areas^15^.

## Methods

We produced maps for species associated with at least one terrestrial habitat in the IUCN Habitat Classification Scheme^11^. We excluded a total of 342 species of mammals and 495 species of birds (6.2% and 4.6% out of 5,481 and 10,651 species, respectively). These comprised 135 mammals and 168 birds exclusively associated with marine habitats (i.e., marine neritic, marine oceanic, marine deep ocean floor, marine intertidal or marine coastal/supratidal), 29 mammals exclusively associated with caves and subterranean habitats, 131 mammals with no associated habitat codes, 8 mammals and 162 birds classified as Extinct, 1 mammal and 5 birds classified as Extinct in the will, 12 mammal and 142 bird species that are restricted to small islands not included in the land-cover map we used, and 26 mammals and 18 birds that had null AOH, caused by errors in the coding of habitat and elevation^16^.

Species may have more than one range polygon, coded according to presence (the species is or was in the area), origin (why and how the species is in the area) and seasonality (seasonal presence of the species in the area)^17^. We used as a base for the AOH maps a predetermined subset of the IUCN Red List range ^18^polygons for each species^16^. Following the Global Standard for the Identification of Key Biodiversity Areas Guidelines^19^, we selected range polygons with *extant* and *probably extant* presence; *native, reintroduced*, and *assisted colonisation* origin; and *resident* seasonality for non-migratory species (all mammals and non-migratory birds; 8,979 species). For migratory birds (1,816 species), we kept separate the ranges for *breeding* (1,446 species), *non-breeding* (1,550 species) and a combination of *resident* and *uncertain* (1,290 species) seasonality. We provide an R script to merge the AOH sub-maps into a single composite map for each species. For 18 mammal and 22 bird species classified as Critical Endangered, there were no presence polygons coded as *extant* or *probably extant*. As the conservation of these species is a priority, we produced AOH maps using the *possibly extinct* polygon for these taxa.

AOH maps are produced by subtracting unsuitable areas from range maps, using data on each species’ associated habitat. As habitats in the IUCN Red List are not spatially explicit (although we note the existence of recently published maps^20^), we used a recently published translation table^12^ based on the Copernicus Global Land Service Land Cover (CGLS-LC100)^21,22^ and the European Space Agency Climate Change Initiative land cover 2015 (ESA-CCI)^23^. We developed the AOH maps based on CGLS-LC100 as CGLS-lC100 has a higher resolution and accuracy than ESA-CCI. CGLS-LC100 is in a regular latitude/longitude grid (EPSG:4326) with the ellipsoid WGS 1984 with a grid resolution of 1°/1008 or approximately 100 m at the equator, defining the resolution of the AOH maps. The translation table presented the relation between each habitat in the IUCN Classification Scheme and each land-cover class as a continuous variable. To create a binary table of association or non-association, Lumbierres et al.^12^ proposed three potential thresholds based on the tertiles of the positive association values of the table. We produced maps for the three proposed thresholds and evaluated the ratio of AOH area to range area. As the threshold increased, the ratio decreased, and the results were more similar to previous AOH maps ^6^. Dahal et al.^16^ evaluated these three thresholds and corroborated that an increase in the threshold did not reduce the performance of the AOH maps during validation. Therefore, we present the maps produced using the highest threshold (odds ratio> 1.7). Species’ elevation limits were extracted from the IUCN Red List ^18^. To subtract the parts of the range outside the elevation limits, we used the Shuttle Radar Topography Mission^24^ map, resampled at the resolution of the CGLS-LC100.

One of the main complexities of this analysis was the large amount of data generated in the process. Therefore, the AOH maps were produced using the GRASS GIS^25^ software, which allows processing large amounts of raster data efficiently. The AOH production procedure consisted of four steps, following Rondinini et al. (2011)^6^: 1) Transforming the habitat codes of each species into land cover classes using the translation table^12^. 2) Creating a base map that combines the information on land cover and elevation 3) Creating reclassification files containing the information on land cover and elevation preferences for each species. 4) Reclassifying the base map based on the reclassification files to create the AOH for each species. We also created intermediate AOH maps clipped only by elevation or only by land cover, in order to explore the influence of each of these parameters on the final AOH. Once the AOH were produced, we calculated richness maps by stacking the AOH maps.

### Data Records

With this article we accompany a sample, the full data sate is store in Dryad Open Access Repository. The AOH data, including tables and maps. The data are organised by taxonomic Class and Order with zipped folder by Order. In the case of birds, we separated migratory species from non-migratory species. In each class folder, maps are organised by taxonomic. AOH maps in GeoTiff. An additional folder contains the richness maps for each class of all species and of globally threatened species. In each folder, we include a table with information of the excluded species, indicating the reason for exclusion; and a table with the included species and the AOH/range ratio. For migratory birds, we included a table specifying which maps (breeding, non-breeding and resident) each species has and code to merge the different parts of the AOH.

### Technical Validation

The accuracy of the AOH maps was assessed using a novel methodology developed by Dahal et al. ^16^ and full details of the validation are provided there. This methodology allowed validation of AOH maps for species with or without point localities. Previous AOH maps were validated only using point localities and polygons of occurrence ^6–8^, leaving some of the AOH maps unvalidated.

Our method employed a two-step approach. The first step identified potential systematic errors in the AOH maps using a modelling approach. This approach flagged 178 and 64 AOH maps for birds and mammals respectively that were carefully studied to identify the sources of potential errors. These potential errors were caused by inaccuracies in species’ elevation limits, habitat coding or the translation table^12^ used to assign habitat to land cover. Work is currently underway to address these issues, and improved AOH maps will be available in the future for download at https://www.iucnredlist.org/resources/grid/spatial-data. A complete list of flagged maps can be found in Dahal et al. ^16^

The second step used point localities to validate the maps at the species level. To validate the AOH maps, the proportion of points localities falling inside the AOH (point prevalence) was compared with the A.O.H./range ratio (model prevalence). If point prevalence exceeded model prevalence, the AOH was assumed to be better than a random distribution within the species’ range^6^. This was done for the 4889 birds (46% of all bird species) and 420 mammals (8% of all mammal species) that had available point locality data. For mammals, this represented 157 species more than in a previous set of AOH^6^ maps published in 2011. AOH maps were better than random for 95.9% bird and 95 % mammal species. The unavailability of point locality data for half of bird species and most mammal species remains a major limitation of the validation analysis. However, the first step of the method allowed us to assess at least the general soundness of AOH maps for species that did not have suitable point localities for validation.

### Usage Notes

The maps are presented in raster byte GeoTIFF format. The values of the maps are 1 for the AOH area and Null for the background. The geographical extent of each map is defined by the species’ range. Each species map is presented separately with the species binomial name, and the genus and specific epithet separated by an underscore. For migratory birds we produced three different maps, that are coded using, R, B and N for resident, breeding and non-breeding AOH maps, respectively. We present code written in R to merge the different AOH maps for migratory species according to the needs of the user. For species with null AOH we recommend using the mapped range.

## Code Availability

The code to produce the AOH is derived from code produced by Rondinini et al. (2011)^6^. AOH maps are produced reclassifying a base map that contains information on elevation and land cover. The geographical range maps are used to mask the areas outside the distribution of the species. Each species has a reclassification file that indicates which land cover classes and elevations are suitable. To transform the habitat information into land cover we used the translation table ^12^. The code is both in GRASS and R.

### Base map

The base map is the map that is reclassified to produce the AOH. Each cell value is a combination of land cover and elevation, where the three first digits represent land cover and the three last digits elevation in m/10.

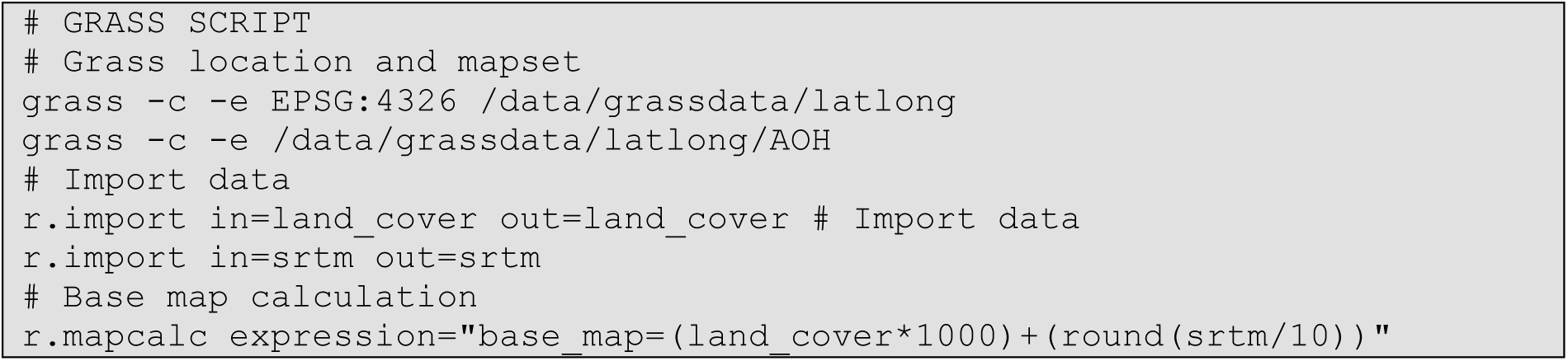

### Reclassification Files

The GRASS reclass function has a specific format for the reclassification instructions. The script produces reclass files to apply to the base map in GRASS to produce maps of area of habitat for terrestrial species. It reads a file that contains land cover associations, with the following column headers: species name, one column per land cover class (with numeric column names for land cover; e.g., 10, 20, 210), and two columns representing elevation range (elevation_min and elevation_max). If the elevation range for a species is unrecorded, it is set to 0-9000 m.

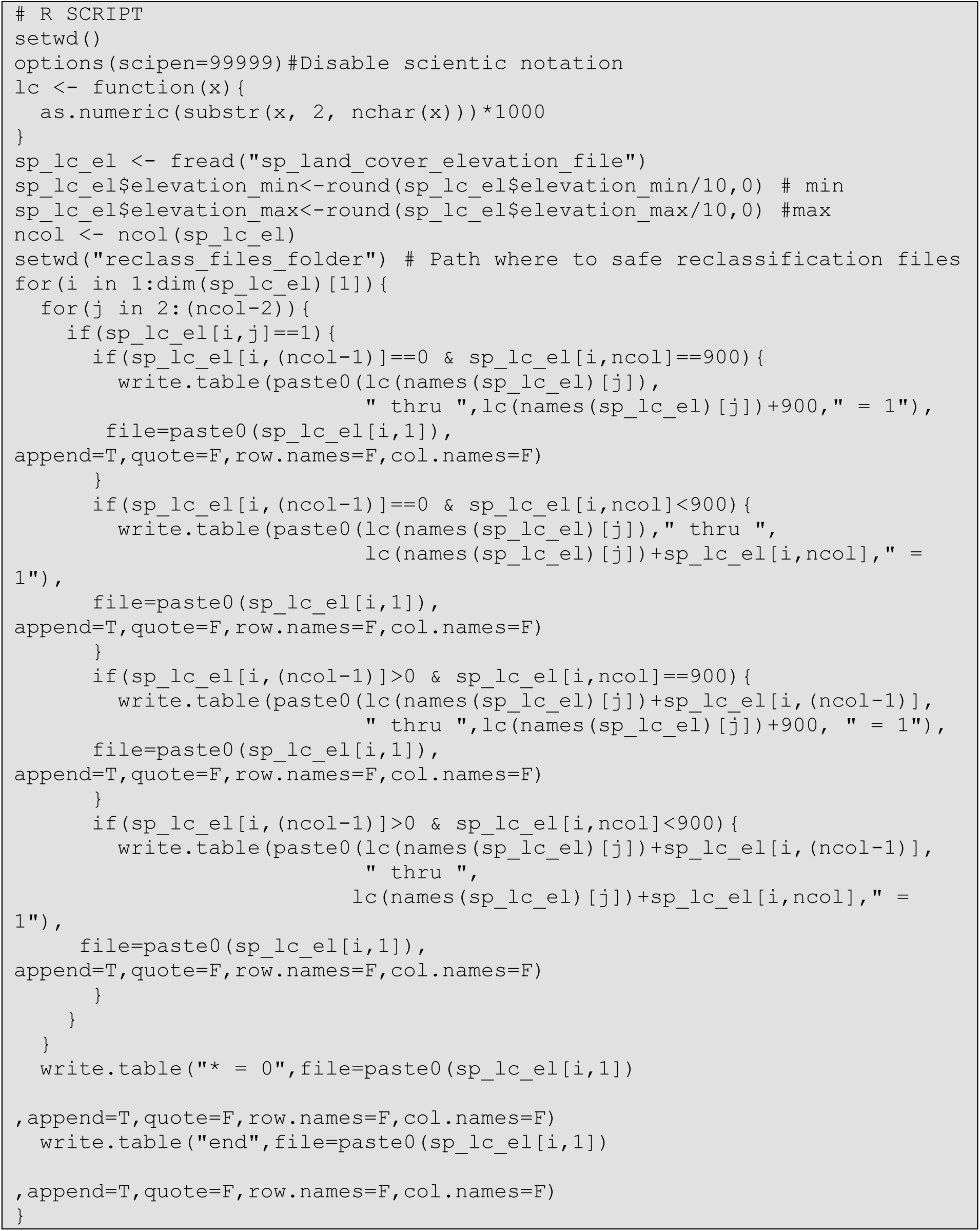

### AOH production

AOH is confined inside the geographical range. The geographical range maps can be downloaded from https://www.iucnredlist.org for mammals, and http://datazone.birdlife.org/species/requestdis for birds. The ranges are imported into GRASS and rasterised. The ranges are used to mask the area outside the species distribution. Inside the non-masked areas, the base map is reclassified using the reclassification file.

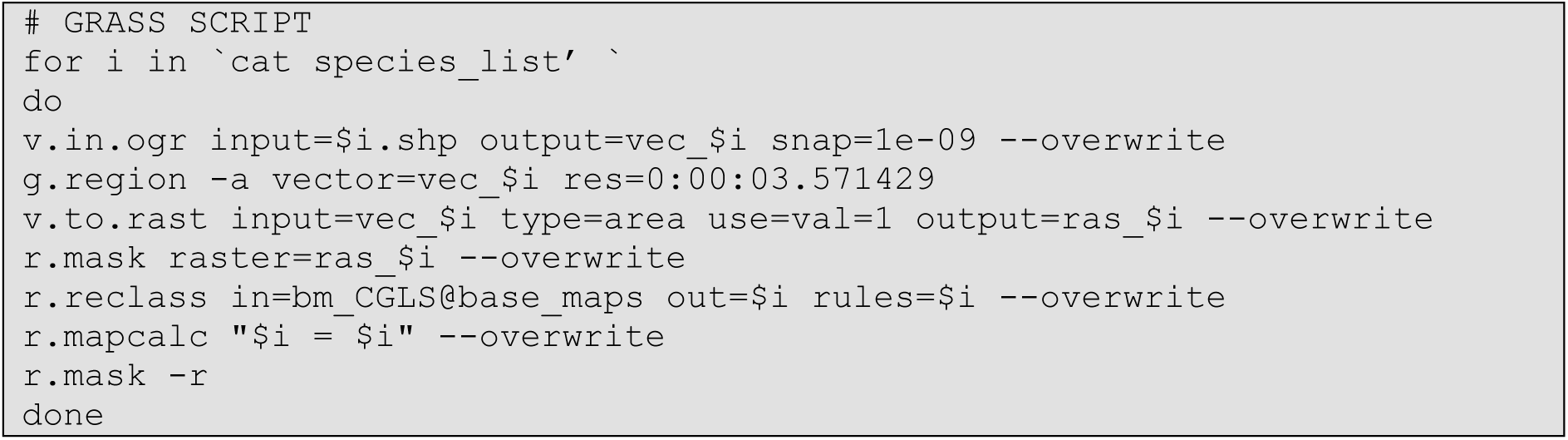

## Acknowledgements

This research is part of the Inspire4Nature Innovative Training Network, funded by the European Union’s Horizon 2020 research and innovation program under the Marie Skłodowska-Curie grant agreement no. 766417. MDM acknowledges support from the M.U.R. Rita Levi Montalcini programme.

## Author contributions

ML, CR, PFD, PRD and SHMB conceived the study. CR and ML developed the code for the analysis. ML, CDS and PRD developed the analysis. ML led the writing of the manuscript. All authors contributed to drafts and gave final approval for publication.

---

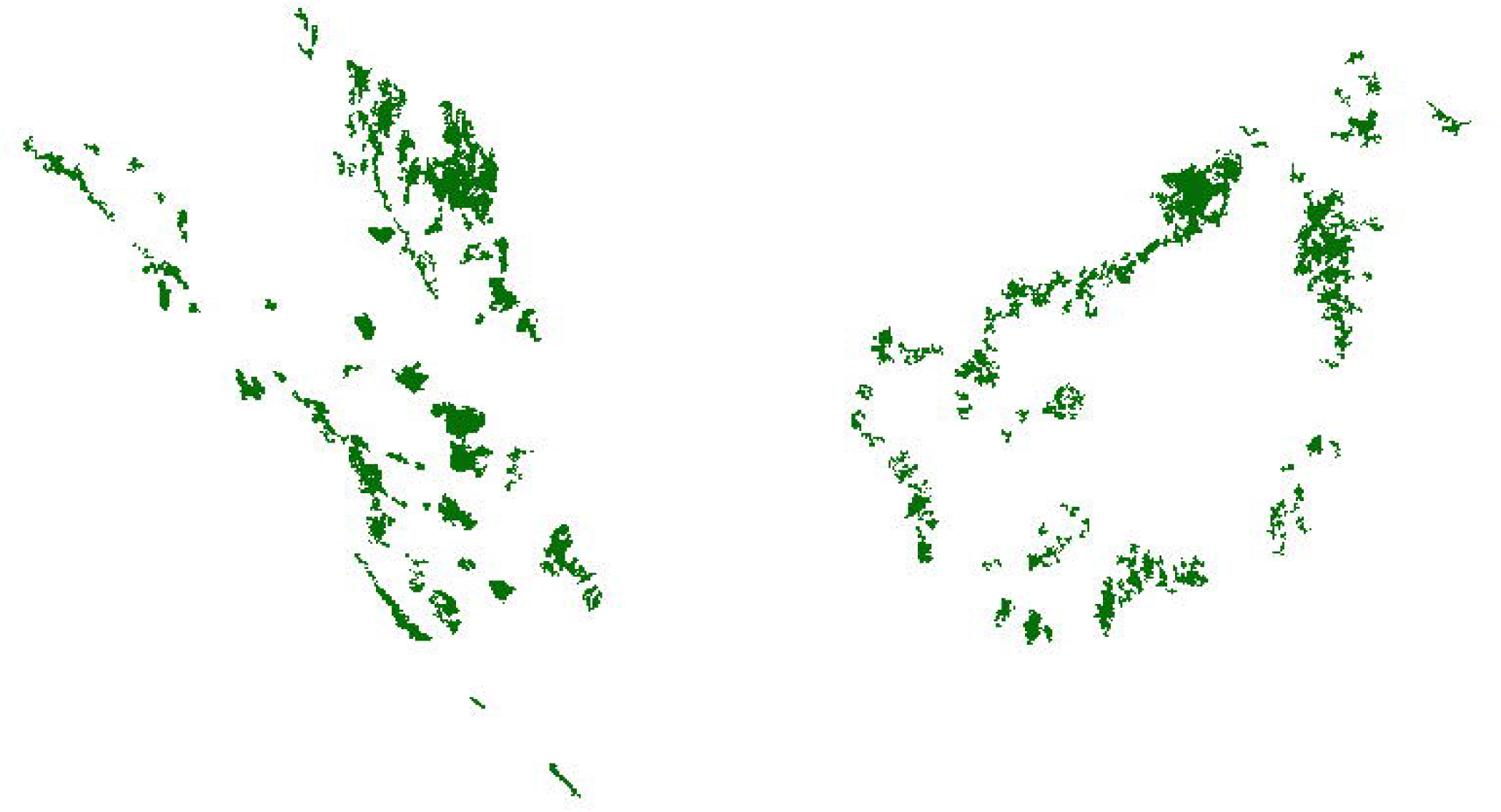

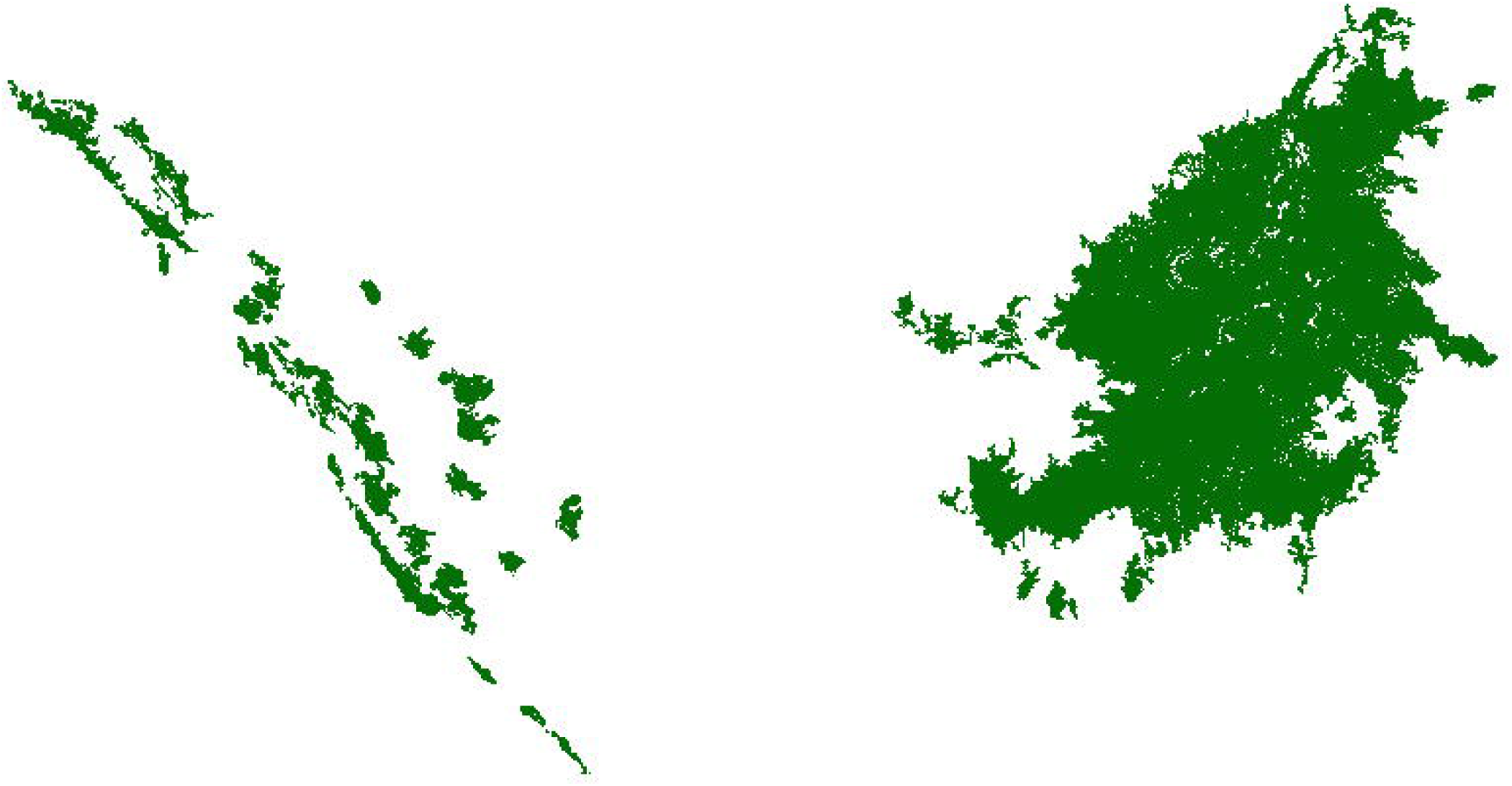

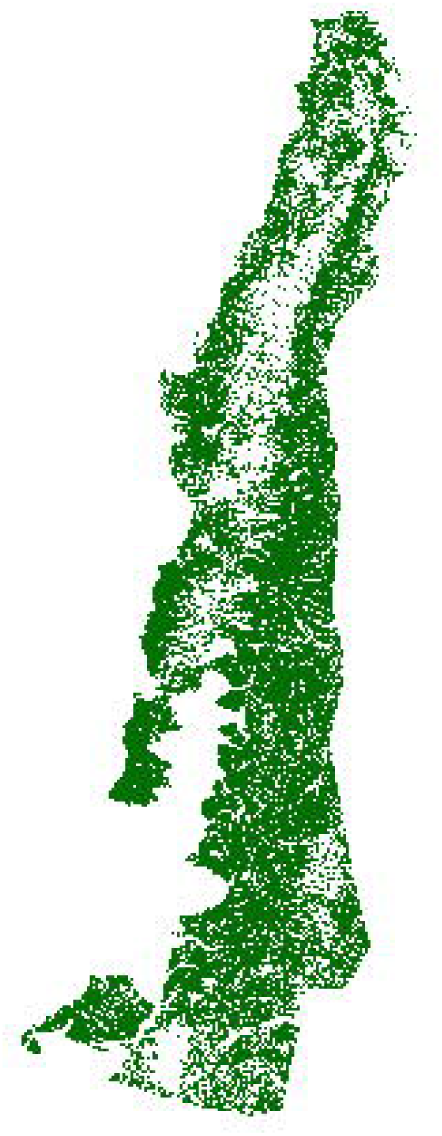

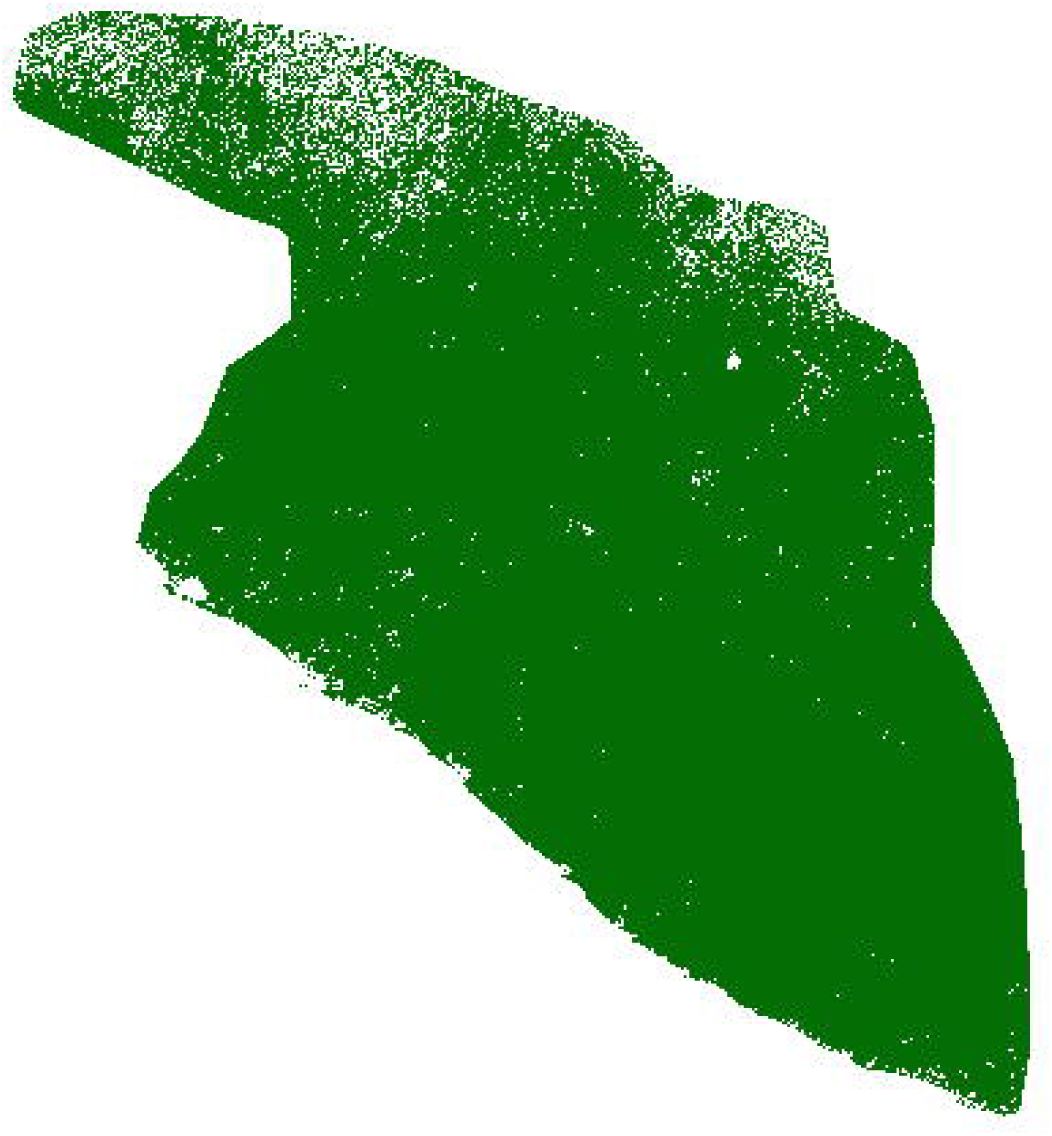

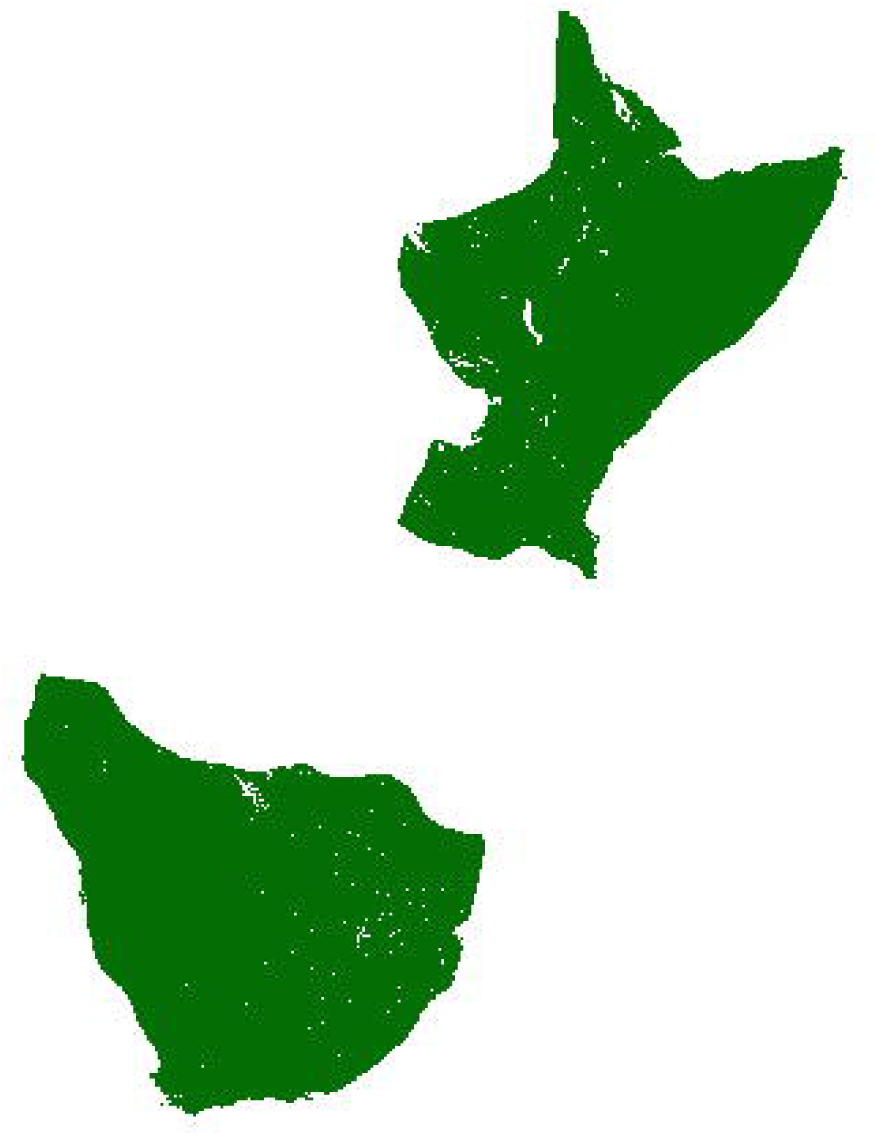

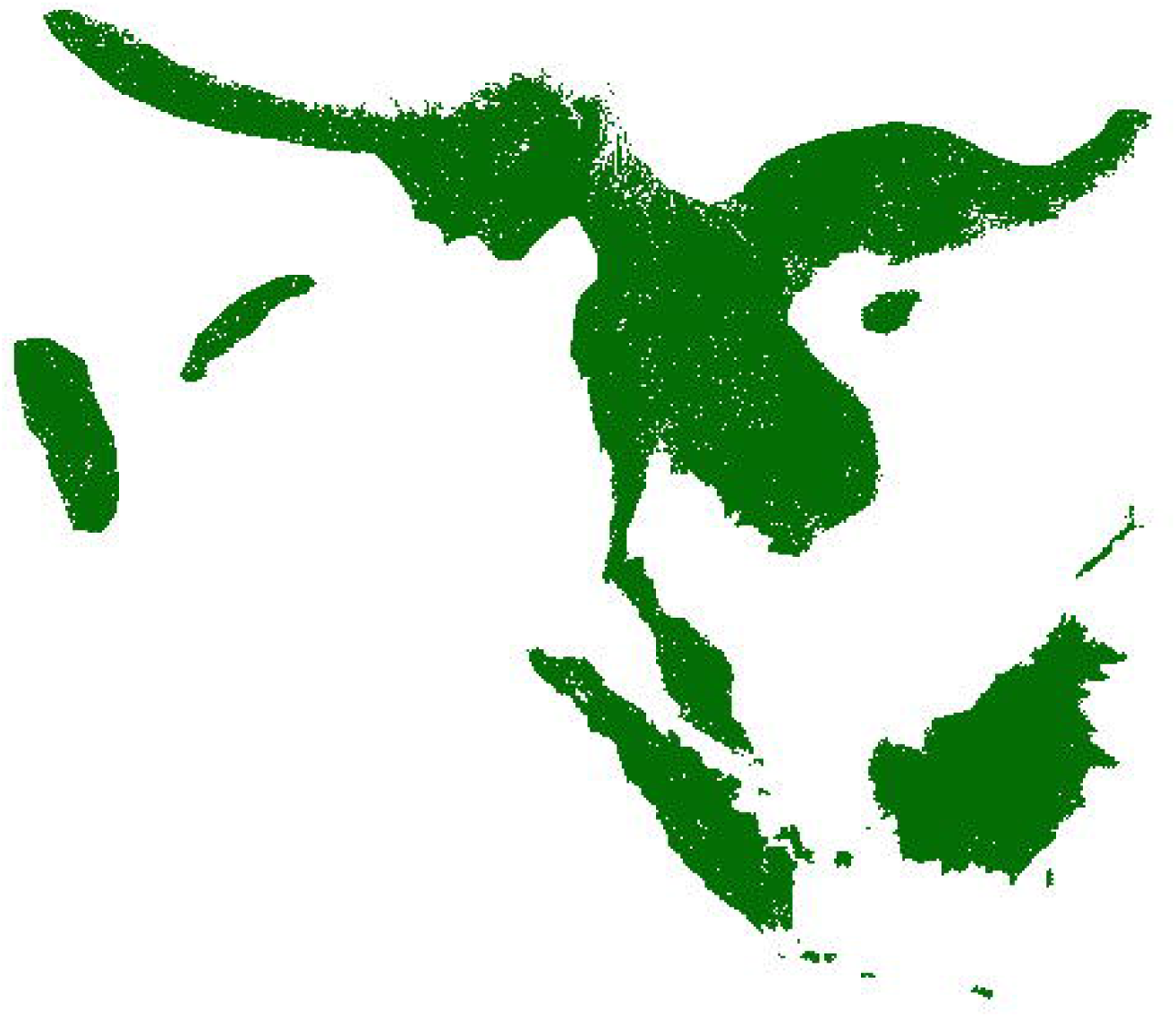

